# Ethylene-induced host responses enhance resistance against the root-parasitic plant *Phelipanche aegyptiaca*

**DOI:** 10.1101/2025.10.05.680554

**Authors:** So-Yon Park, Chong Yang, James H Westwood

## Abstract

The root parasitic plant *Phelipanche aegyptiaca* and related species pose a grand challenge for agriculture. By directly attaching to host crops to acquire resources, they can severely decrease yields. At the same time, their underground location hides them from sight for much of their life and protects them from typical weed control methods. New strategies for parasitic weed control are needed, with host resistance among the most attractive, but the mechanisms by which hosts respond to parasitic plant attacks are still unclear. In plants, the phytohormone ethylene is a crucial modulator responding to various stresses such as flooding and pathogen attack, and our data suggest that ethylene signaling may be important in host response to *P. aegyptiaca*. Here, we demonstrated that ethylene plays a role against *P. aegyptiaca* using two host plants, *Arabidopsis* and tomato. *Arabidopsis* plants with the ethylene reporter construct (*EBS::GUS*) were analyzed and revealed that stress from excess water and *P. aegyptiaca* parasitism both induced ethylene responses in the host *Arabidopsis* roots. We also observed that applying an ethylene precursor (ACC) to host roots inhibits the attachment of *P. aegyptiaca*. Lines of *Arabidopsis* and tomato with mutations in *Ethylene-Resistant 1* (*ETR1*) and *Constitutive Triple-Response 1* (*CTR1*) have altered tolerance to *P. aegyptiaca*, suggesting that the ethylene signaling pathway is associated with enhanced resistance to parasitism. These results point to ethylene-mediated responses as a starting point to gain insight into host response to parasitism, with potential to increase host resistance to parasitic plants.

## Introduction

*Phelipanche* and *Orobanche* species (broomrapes) are non-photosynthetic, obligate parasitic plants with a highly modified life cycle. They attack the roots of compatible host plants, and their development is tightly linked to their hosts, beginning with the need for a specific host-derived signal to trigger germination and continuing through flowering (Yoneyama *et al*., 2013). Parasitic plants connect to their hosts via specialized structures, termed haustoria, that function to withdraw water and nutrients from the hosts (Westwood *et al*., 2010). The interaction between parasitic plants and their hosts provides one of the most direct examples of plant-plant interactions, in which one species invades another, manipulating the defenses and nutrient allocation of the host to create an apparently seamless unity (Clarke *et al*., 2019). The mechanisms by which this is accomplished are poorly understood.

Parasitic plants of the family Orobanchaceae include some of the most destructive parasitic weed species, including the broomrapes and witchweeds (*Striga* spp.) (Nickrent & Musselman, 2010). Together, these parasites impact crop production in Africa, the Mediterranean, and Europe, with infestations spreading throughout the world (Casadesús & Munné-Bosch, 2021; Vurro, 2023). The broomrapes specifically affect dicotyledonous crops, including economically important tomato, sunflower, legumes, and many more (Aly & Dubey, 2014). These parasites are exceptionally challenging to detect and control because they spend their vegetative life underground, attached to host roots, and inflict most of their damage to the crop before they emerge from the soil. An ideal solution to the problem posed by these parasites is to develop host crops that are resistant to attack. Such technology would be simple to employ, environmentally benign (no spraying of chemicals), and inexpensive. The challenge is that instances of host resistance to parasitic weeds are relatively rare, and only a few of these have been characterized. A more complete understanding of host-parasite interaction is needed to understand which processes, pathways, and genes are most promising for manipulation to tip the interaction in favor of host resistance.

Ethylene has long been known as a critical regulator of plant development but is increasingly recognized as playing essential roles in various biotic and abiotic stresses (Groen & Whiteman, 2014; Broekgaarden *et al*., 2015). Ethylene consists of two main aspects, the biosynthesis pathway and the signaling pathway (Ju & Chang, 2015). Ethylene is synthesized from methionine through a three-step pathway that is subject to feedback regulation. First, methionine is converted to AdoMet and 1-aminocyclopropane-1-carboxylic acid (ACC) by S-adenosylmethionine synthesis (SAMS) and ACC synthase (ACS). ACC is then oxidized to ethylene by ACC oxidase (ACO) (Wang *et al*., 2002). The transcriptional and post-translational modification of ACS regulates cellular ACS activity, which is a critical element in fine-tuning ethylene production (Xu & Zhang, 2015). Once ethylene is synthesized, its effects are controlled by genes of the signaling pathway, such as *Ethylene-Resistant* (*ETR*), *Constitutive Triple-Response* (*CTR*), and *Ethylene-Insensitive* (*EIN*).

The ethylene signaling pathway in plants functions in early response to the presence of invaders such as fungal, bacterial, and insect pathogens (van Loon *et al*., 2006; Broekgaarden *et al*., 2015; Xu & Zhang, 2015). After pathogens trigger ethylene signaling, transcription factors of the EIN family play a dominant role in the activation/suppression of downstream pathway genes related to plant defense mechanisms. For example, plants with *ein2* and *ein3* knockout mutations are susceptible to many fungal and bacterial infections (van Loon *et al*., 2006). On the contrary, plants overexpressing *Ethylene Response Factor 2* (*ERF2*) present enhanced resistance to pathogens and insects (Pre *et al*., 2008; Anderson *et al*., 2010; Zhu *et al*., 2013; Broekgaarden *et al*., 2015).

Ethylene has been studied in parasitic plants of the Orobanchaceae. For example, the facultative parasites *Triphysaria versicolor* and *Phtheirospermum japonicum* appear to use ethylene as a pivotal hormone to control haustorium growth (Tomilov *et al*., 2005; Cui *et al*., 2020). However, the host ethylene response has not been well studied in host-parasite interactions. In this study, we observed the host ethylene pathway against the model root parasitic plant, *Phelipanche aegyptiaca*, which is well suited to laboratory studies and readily parasitizes the model plant *Arabidopsis thaliana* (Westwood, 2000). We found that *P. aegyptiaca* induces the ethylene signaling pathway in the host and is correlated with suppression of tubercle growth. We suggest that the ethylene pathway in host plants holds promise for identifying mechanisms to enhance protection against *P. aegyptiaca*.

## Material and Methods

### Plant materials and growth conditions

*Arabidopsis* and tomato seeds were stratified in water at 4 °C for a day and then sown onto Sungro Professional Growing Mix. Plants were grown in a Conviron (Controlled Environments Inc) growth chamber with 23 °C, 9-hour light, and 15-hour dark cycles for 2 weeks. Fifteen days old plants (*Arabidopsis* and tomato) were removed from the soil, transferred to glass microfiber filter (GFA) paper (Whatman) in polyethylene (PE) bags (Rubiales *et al*., 2006). Hydroponic nutrient solution (Hoagland’s solution) (Hoagland & Arnon, 1938) was applied to the PE bag during experiments. 26 × 9 cm PE bags were used to hold multiple *Arabidopsis* plants, and 26 × 40 cm PE bags were used for each tomato plant.

### *Arabidopsis* and tomato mutants

*Arabidopsis EIN3 binding site* (*EBS*) promoter*::GUS* was from Stepanova lab, NCSU (Stepanova *et al*., 2007). *Arabidopsis etr1-1* (CS237) and *ctr1-1* (CS8057) were distributed from The *Arabidopsis* Information Resource (TAIR) (arabidopsis.org). Tomato mutant lines, *etr1* (LA4477, *nr*) and *ctr1* (LA4484, *epi*), were obtained from the Tomato Genetics Resource Center (TGRC) (https://tgrc.ucdavis.edu/). *Arabidopsis* Col-0 and tomato var. Micro-Tom were used as control plants.

### *P. aegyptiaca* inoculation and quantification

*P. aegyptiaca* seeds were conditioned and inoculated according to our previous methods (Clermont *et al*., 2019; Clarke *et al*., 2020). Using a fine-tipped paintbrush, individual *P. aegyptiaca* seeds were placed onto roots of 4-week old host (*Arabidopsis* or tomato) (Figure 1A, black box). The total numbers of *P. aegyptiaca* tubercles on host roots were counted using a dissecting microscope (Zeiss Stemi SV11) 2 weeks after inoculation. The attached *P. aegyptiaca* tubercles were further classified as either initial attachment, early-stage tubercle, or late-stage tubercle, which correspond to the stages 3, 4.1, and 4.2 in the Parasitic Plant Genome Project datasets (Westwood *et al*., 2012). At least three independent experiments were conducted. Statistical analysis was carried out using JMP (JMP Software, USA). One-way ANOVA followed by Tukey’s multiple comparisons was the method used for significance tests.

**Figure 1.**
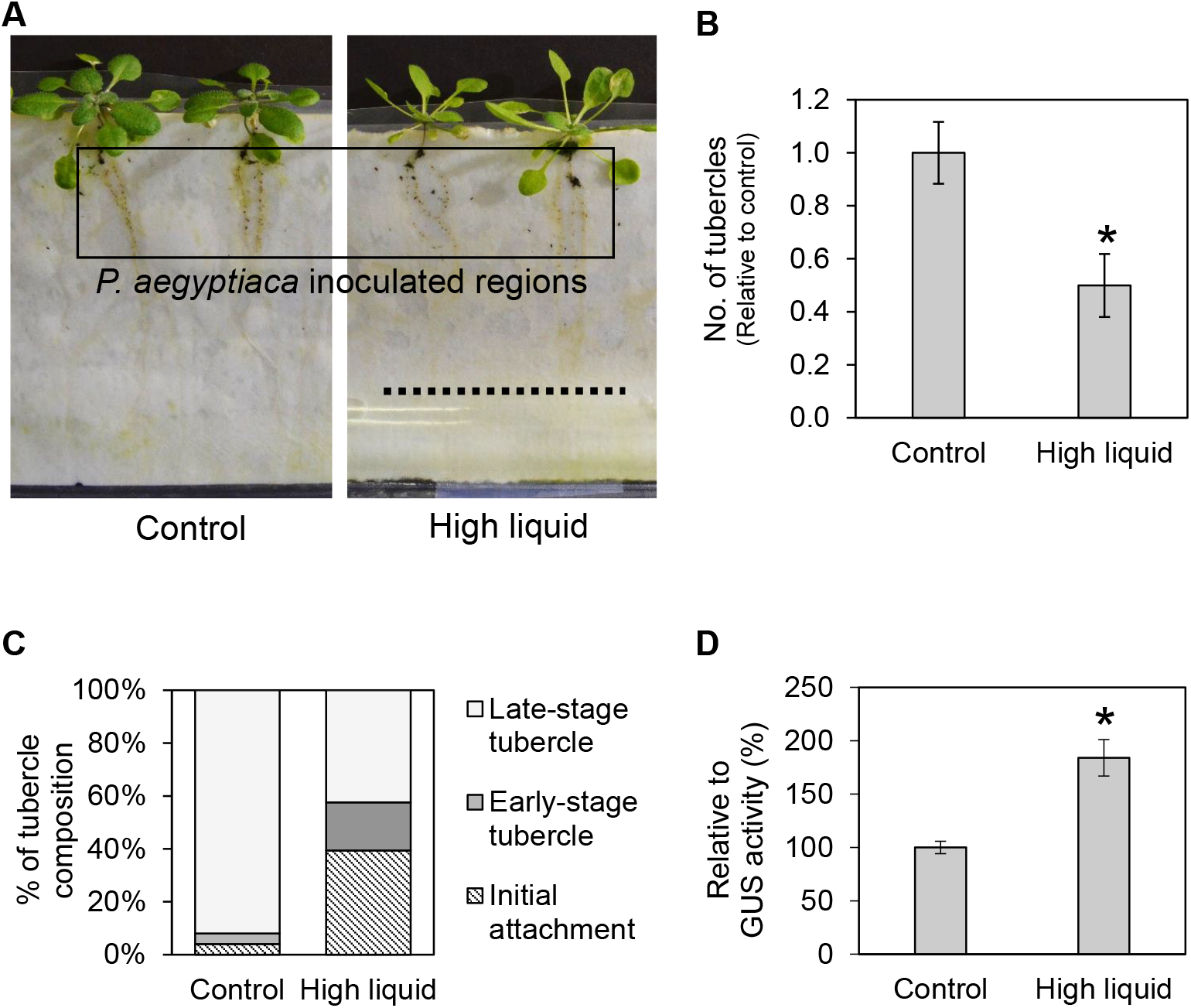
*Phelipanche aegyptiaca* inoculation assay and different liquid levels. **A**. *P. aegyptiaca* were inoculated on 4 weeks old *Arabidopsis* plants. High liquid treatment (dashed line indicates upper limit of solution) started when parasites were applied to the host roots (black box). **B**. Numbers of parasites were counted 14 days after *P. aegyptiaca* inoculation (DAI). Data are means with SE indicated. **C**. % of tubercle composition under control and high-liquid condition. **D**. Relative GUS activity with quantitative MUG assay. GUS activity (pmol MU/min/mg protein) was measured in *EBS::GUS* after high liquid treatment. Means and SE from 3 independent experiments are shown. The bar marked with * is different from the control at P < 0.05 (ANOVA, Tukey’s test).

### High liquid treatment under *P. aegyptiaca* infection

For the ideal growth condition, PE bags with *Arabidopsis* needed around 3-5 ml of hydroponic nutrient solution every day to maintain the humidity inside the bags without extra liquid around. Four bags (4 Col-0 plants/bag) were grown under the ideal liquid level until *P. aegyptiaca* infection. Forty seeds of germinated *P. aegyptiaca* were applied to the roots of each host plant. After *P. aegyptiaca* inoculation, we randomly selected 2 bags to receive optimal levels of liquid (control), and another 2 bags to receive excess liquid (treated). For the simulated flooded treatment, 15-20 ml of the hydroponic nutrient solution was maintained such that the hydroponic solution reached the black dashed line in Figure 1A. The flooding condition lasted for 2 weeks. Two weeks post-parasitism, the total number of tubercles growing on *Arabidopsis* roots were counted and classified by developmental stage. Results were obtained from at least three independent biological replicates.

### Quantitative GUS Assays

The fluorescent β galactosidase assay with 4-methylumbelliferyl ß-D-glucuronide (4-MUG) (Fisher) was conducted to detect the ethylene response under the high liquid treatment using *Arabidopsis* plants with a GUS reporter system, *EBS::GUS* (Stepanova *et al*., 2007). Briefly, 15-day-old *EBS::GUS Arabidopsis* plants were transplanted from soil/pots to PE bags (4 *plants per* bag). The *Arabidopsis* plants were exposed to control or excess liquid conditions for 2 more weeks, and then GUS activity was analyzed by the 4-MUG assay. Total proteins were extracted by the protein extraction buffer according to the manufacturer’s guidelines (#AAB2107503, Fisher). Concentrations of total proteins were quantified by Bradford assay (Bio-Rad) using bovine serum albumin (Promega) as the standard. For the 4-MUG assay, fluorescence was detected at the excitation/emission wavelengths of 365 nm / 455 nm using a plate reader (Biotek Synergy HT). A standard curve was generated from 0.1 to 50 μM of 4-methylumbelliferone (4-MU) (Sigma). The GUS enzyme activity was expressed as picomoles of 4-MU produced per milligram protein per minute (pmol MU/min/mg protein). Data were generated from three independent biological replicates.

### Histochemical GUS Assays

For the histochemical GUS staining, 10-day-old *P. aegyptiaca* with *Arabidopsis* Col-0 and *EBS::GUS* (Stepanova *et al*., 2007) roots were stained by 5-bromo-4-chloro-3-indolyl-BD-glucuronide (X-gluc) for 3 hours following the manufacturer’s guidelines (Sigma). Destained tissues were photographed using a stereo-zoom microscope (Discovery V12, Carl Zeiss).

### ACC treatment

Ethylene precursor 1-aminocyclopropane-1 carboxylic acid (ACC) (Sigma) was used to induce ethylene responses in plants (Van de Poel & Van Der Straeten, 2014). 14-day-old *Arabidopsis* wild-type (Col-0) plants were transferred from soil and grown in PE bags as discrived above. 100 μM ACC mixed in hydroponic nutrient solution was applied to the roots from the day of *P. aegyptiaca* inoculation (0 day after inoculation, 0 DAI) to 14 DAI or from 5 DAI to 14 DAI. On day-14 after inoculation, the total number of *P. aegyptiaca* tubercles were counted. For the negative control, only DMSO mixed with the hydroponic nutrient solution was applied to roots. One-way ANOVA followed by Tukey’s multiple comparisons were used to test for significant differences. At least three independent biological replicates were analyzed for each treatment.

## Results

### High liquid conditions inhibit the attachment and growth of *P. aegyptiaca*

Growing *P. aegyptiaca* in a polyethylene (PE) bag growth system demands precision when applying the hydroponic nutrient solution. Too little solution may lead to drought stress, while too much inhibits tubercle formation. The reason for inhibited tubercle success under high moisture conditions has not been elucidated, but since an inhibitory level of liquid is well below the level of parasite seeds (Figure 1A), the effect does not seem to be due to submersion of seeds. To study the inhibition of *P. aegyptiaca* under conditions of excess moisture, we first compared the number of tubercles on host roots under optimum liquid conditions (control) and high liquid conditions. The numbers of tubercles formed under the high liquid treatment were 50% lower than the number of tubercles formed under control conditions (Figure 1B). For a more detailed analysis, the *P. aegyptiaca* tubercles were classified into development stages. Under control conditions, the great majority (92%) of *P. aegyptiaca* tubercles developed to the late-tubercle stage (Figure 1C). However, under high liquid conditions only 42.4% of *P. aegyptiaca* tubercles were in the late-tubercle stage, and 39.4% were still in the initial attachment stage.

Flooding conditions are known to induce ethylene response in plants (Grichko & Glick, 2001; Steffens, 2014). To investigate whether the high liquid treatment increases the ethylene response in host plants, we used an *Arabidopsis* ethylene response GUS reporter system, *EIN3 binding site* (*EBS*) promoter*::GUS* (Stepanova *et al*., 2007). This *EBS::GUS* plant expresses GUS activity in roots and shoots in response to ethylene and can be measured using the fluorescent β-galactosidase assay with 4-MUG. The *EBS::GUS* reporter plants exposed to control conditions had 63 pmol MU/min/mg protein, while the high liquid condition had 116 pmol MU/min/mg protein (Figure 1D), a 184% increase over the control.

### *P. aegyptiaca* increases the ethylene response of the host

To determine whether parasitism by *P. aegyptiaca* affects the ethylene response of host roots, the same *EBS::GUS* transgenic *Arabidopsis* lines described above were challenged with the parasite. For this experiment, *EBS::GUS* and wild-type *Arabidopsis* Col-0 were grown under ideal growing conditions in PE bags (i.e., without excess water). To observe expression of the ethylene-responsive GUS reporter system under parasitism, *P. aegyptiaca* infected host roots were stained 10 days after parasite attachment. While no GUS activity was detected in Col-0 roots (black arrows, Figure 2A), *EBS::GUS* roots had blue color indicating GUS enzyme activity in the region of parasite attachment (black arrows, Figure 2B and 2C).

**Figure 2.**
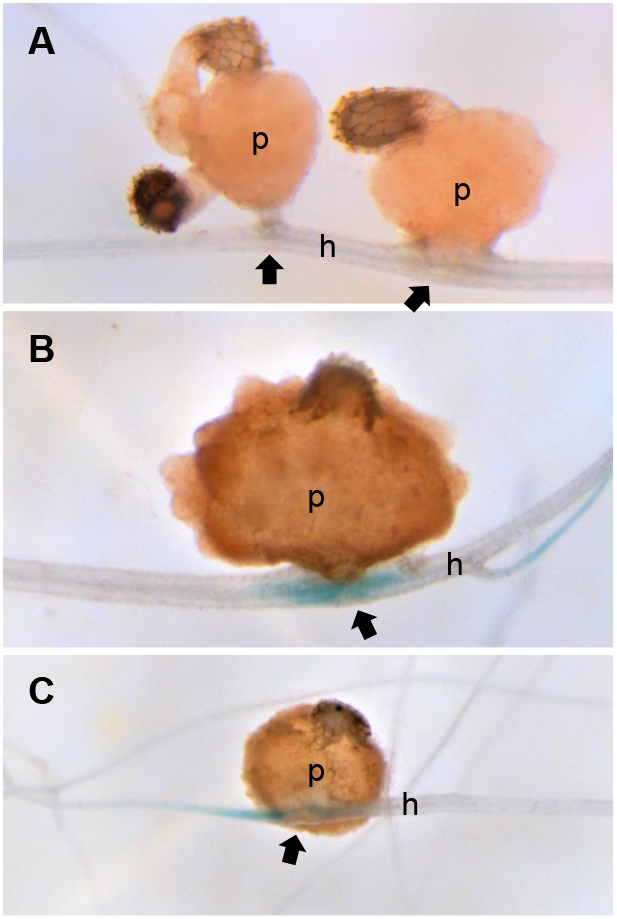
Expression patterns of *EBS::GUS* ethylene reporter in roots with *P. aegyptiaca*. Histochemical staining of GUS activity was performed 10 days after *P. aegyptiaca* inoculation in *Arabidopsis* Col-0 (**A**) and *EBS::GUS* ethylene reporter (**B-C**) transgenic plants. p = *P. aegyptiaca*. h = host root. Arrows indicate a region of parasite attachments.

### The ethylene precursor ACC suppresses attachments of *P. aegyptiaca*

To investigate the effect of induced ethylene synthesis on parasitism, roots of *Arabidopsis* Col-0 that were inoculated with *P. aegyptiaca* were treated with ACC. As an ethylene precursor, ACC is a plant growth regulator alternative to ethylene and is widely used to induce ethylene responses when ethylene or ethephon is unavailable (Zhang *et al*., 2010). ACC applications started either from 0 DAI to 14 DAI or 5 DAI to 14 DAI. Counting the number of tubercles 14 DAI revealed a significant reduction in the number of parasites on ACC-treated plants (Figure 3). Application of ACC on 5 - 14 DAI decreased the number of tubercles by 55%, and when the ACC was applied on 0 - 14 DAI, the reduction was 80% compared to mock treated (DMSO) controls.

**Figure 3.**
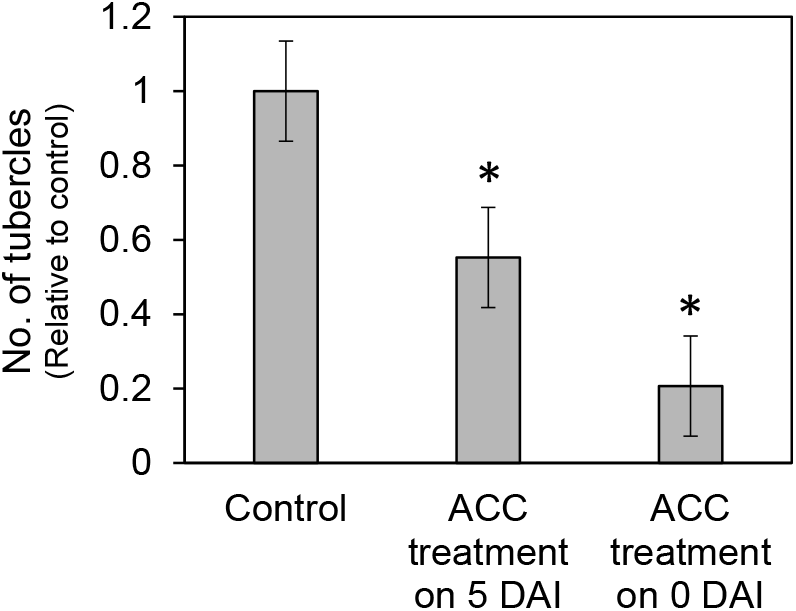
Effect of ACC treatment on *P. aegyptiaca* attachment rates. *Arabidopsis* Col-0 were inoculated with *P. aegyptiaca* and 100 μM ACC (dissolved in DMSO) was applied either on the same day (0 DAI) or 5 DAI. Control was treated with DMSO only. Data are means of at least 14 replicates with SE indicated. Bars marked with * are different from the control at P < 0.05.

### Hosts with mutations in ethylene signaling show altered response to *P. aegyptiaca*

*Arabidopsis* and tomato plants with mutations in the ethylene signaling pathway were inoculated with *P. aegyptiaca* to investigate how the host ethylene signaling pathway affects parasite success. Both host species have lines mutated in the constitutive ethylene activation gene (*constitutive triple response 1-1*; *ctr1-1*) and the ethylene-insensitive gene (*ethylene receptor 1-1*; *etr1-1*). CTR1 is a negative regulator in ethylene signaling pathway (Lee *et al*., 2017), while ETR1 functions as a key player in the perception of ethylene and its downstream signal transmission (Kugele *et al*., 2022). Compared to control plants, *Arabidopsis* Col-0 and tomato var. Micro-Tom, the number of tubercles on *ctr1-1* plants was significantly lower (Figure 4). In contrast, the *etr1-1* mutants that have reduced response to ethylene supported significantly more parasites than controls.

**Figure 4.**
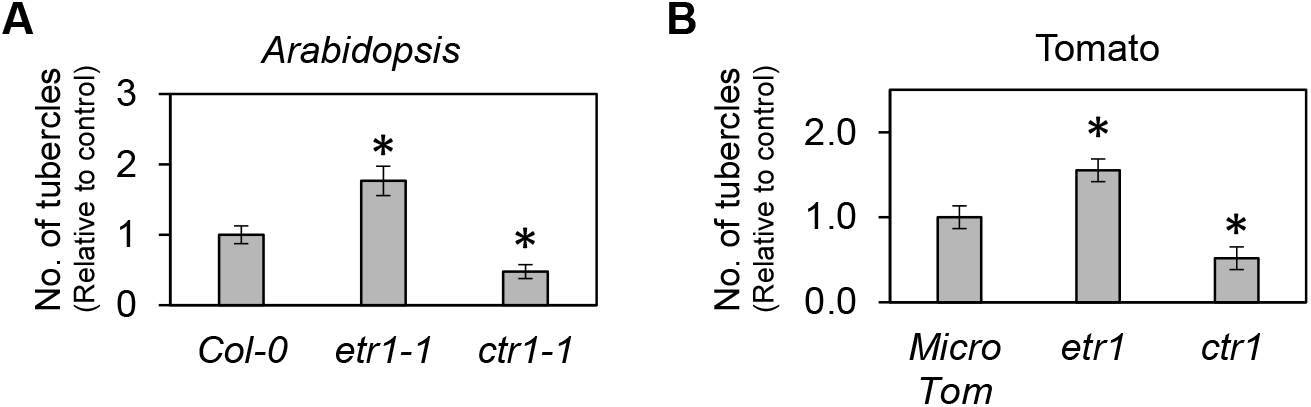
Resistance/susceptibility of ethylene mutant hosts to *P. aegyptiaca. Arabidopsis* plants (**A**) and tomato plants (**B**) were inoculated by *P. aegyptiaca*. The numbers of tubercles were counted 14 days after inoculation. Data are means with SE indicated. Bars marked with * are different from the control at P < 0.05 (ANOVA, Tukey’s test).

## Discussion

Ethylene is recognized as a crucial player in plant-pathogen or plant-insect interactions (Fudali *et al*., 2012; Hu *et al*., 2017; Dubois *et al*., 2018). However, only a few studies have shown the role of ethylene in the host-parasitic plant. For example, ethylene stimulates germination of some root parasites such as *Orobanche ramosa* (Syn. *P. ramosa*) and *Striga* species (Berner *et al*., 1999; Zehhar *et al*., 2002; Zwanenburg *et al*., 2009). The ethylene pathway and other phytohormone pathways are involved in host defense mechanisms against *O. ramosa* and *P. aegyptiaca* (Dos Santos *et al*., 2003; Clarke *et al*., 2020). Ethylene production was associated with early haustorium development in *Triphysaria versicolor*, a facultative parasite in the Orobanchaceae family (Tomilov *et al*., 2005). Similarly, ethylene levels appear to function in seed germination, haustorium growth termination, and host invasion of haustoria in *Phtheirospermum japonicum* (Cui *et al*., 2020).

Unlike previous ethylene-root parasite studies, the work described here focused on the host response to *P. aegyptiaca* attack. It was initiated by the observation that excess liquid in a PE bag environment negatively affected the attachment of *P. aegyptiaca*, and this was confirmed experimentally (Figure 1B). Excess liquid also affected *P. aegyptiaca* growth and development as revealed by the decreased number of late-stage tuberclesas compared to control conditions (Figure 1C). It is possible that high levels of liquid disrupted other aspects of plant signaling, but the finding that these conditions lead to induction of ethylene-responsive genes (Figure 1D), suggests that a stress-related mechanism may underlie these observations. It is possible that enhanced defenses due to the induced ethylene response by the excess water stress (Grichko & Glick, 2001; Steffens, 2014) contributes to suppressed and delayed growth of *P. aegyptiaca*. It is also interesting that the host ethylene response was promoted by *P. aegyptiaca* infection (Figure 2), similar to its induction in plant-pathogen and plant-nematode interactions (Broekaert *et al*., 2006; Gutierrez *et al*., 2009).

To this point the findings were largely correlative, without a direct link between ethylene and *P. aegyptiaca* success. Therefore, to directly induce ethylene responses, we added ACC, which is a direct precursor of ethylene and is used to modulate the ethylene signaling pathway (Broekgaarden *et al*., 2015; Xu & Zhang, 2015). We found that the application of ACC decreased the attachment rate of *P. aegyptiaca* (Figure 3). Additionally, ACC application at 0 DAI was more effective than ACC application from 5 DAI, suggesting that early activation of host ethylene responses was important in suppressing the initial stage of *P. aegyptiaca* attachment. One confounding factor in this experiment is the difficulty in treating only the host plant, without also exposing the parasite to ACC. It is possible that the ACC treatments directly inhibited the *P. aegyptiaca* growth rather than working through activation of host defense mechanisms. To address this issue, we inoculated *P. aegyptiaca* on lines of *Arabidopsis* and tomato having mutations in ethylene signaling. These assays showed that host tolerance to *P. aegyptiaca* was affected by the ethylene response capacities of the hosts (Figure 4). Thus, we conclude that ethylene-mediated host responses can increase the tolerance of host plants to parasitic plants.

These findings build upon previously reported work in which we showed that *Arabidopsis* expressing *35S::ERF2* (e*thylene response factor 2*) was tolerant to *P. aegyptiaca*, and transcript levels of *Arabidopsis ACS6* were up-regulated in *P. aegyptiaca* parasitized host roots (Clarke *et al*., 2020). The ERF2 protein is one of the transcription factors of APETALA2/ethylene response factor family (Sakuma *et al*., 2002) and is involved in biotic/abiotic stress pathways (Nakano *et al*., 2006). ERF2 is also an essential positive regulator for inducing tobacco *ACS* transcripts and ethylene production (2009).

Another prior study provided further support for our hypothesis by demonstrating increased expression levels of ethylene-related genes in host plants infected with *O. ramosa*. (Dos Santos *et al*., 2003). The study reported that transcript levels of *Arabidopsis ACS2* were significantly induced in host roots after placing *O. ramosa* seedlings, indicating that *Arabidopsis* plants activated the ethylene synthesis pathway in response to *O. ramosa* parasitism.

Ethylene also increases the host tolerance against a stem parasitic plant. Ethylene has been reported as a critical defense factor against stem parasite *Cuscuta reflexa* (Hegenauer *et al*., 2016) because a *C. reflexa* tolerant tomato line produced more ethylene than other tomato plants (Hegenauer *et al*., 2016). These reports are consistent with our hypothesis that increased host ethylene response negatively affects the growth of *P. aegyptiaca*.

However, there are conflicting results have been reported as well. *C. campestris* requires host-derived ethylene to extend its searching hyphae into an *Arabidopsis* stem (Narukawa *et al*., 2020). Similarly, host-produced ethylene contributes to *Phtheirospermum japonicum* invasion (Cui *et al*., 2020). The fine-tuning of host-derived ethylene synthesis or signaling pathway may be required for the host-parasitic plant interaction (Narukawa *et al*., 2020). It seems likely that both host and parasitic plants use ethylene in growth and defense, and that signaling may have evolved differently in different types of parasite. In summary, we have shown that elevated water stress and associated ethylene pathway induction are correlated with *Arabidopsis* resistance to *P. aegyptiaca*. Disruption of ethylene signaling in host plants supports a role for ethylene-related defense mechanisms in parasite resistance. This is a promising starting point to further explore the role of ethylene in effective host defense mechanisms.

## Acknowledgements

This work is supported by the Parasitic Plant Genome Project (PPGP) (NSF IOS-1238057), USDA-AFRI-2023-67013-39896 to SP and JHW, NIFA award 160111 to JHW, and National Research Foundation of Korea (NRF) grant funded by the Korea government (MSIT) (No. RS-2024-00336161) to SP. Dr. Christopher R. Clarke at ARS, USDA provided comments for this project. Tomato Genetics Resource Center at University of California, Davis shared tomato mutant seeds. Drs. Anna Stepanova and Jose Alonso (NCSU) provided *Arabidopsis EBS::GUS* seeds.

## Author Contributions

Conceptualization, SP and JW.; methodology, SP, CY, and JW.; writing, SP and JW. All authors have read and agreed to the published version of this manuscript.

## Data Availability

The datasets used and/or analysed during the current study are available from the corresponding authors on reasonable request.

